# Eliminating yellow fever epidemics in Africa: vaccine demand forecast and impact modelling

**DOI:** 10.1101/468629

**Authors:** Kévin Jean, Arran Hamlet, Justus Benzler, Laurence Cibrelus, Katy A. M. Gaythorpe, Amadou Sall, Neil M. Ferguson, Tini Garske

## Abstract

**Background:** To counter the increasing global risk of Yellow fever (YF), the World Health Organisation initiated the Eliminate Yellow fever Epidemics (EYE) strategy. Estimating YF burden, as well as vaccine impact, while accounting for the features of urban YF transmission such as indirect benefits of vaccination, is key to informing this strategy.

**Methods and Findings:** We developed two model variants to estimate YF burden in sub-Saharan Africa, assuming all infections stem from either the sylvatic or the urban cycle of the disease. Both relied on an ecological niche model fitted to the local presence of any YF reported event in 34 African countries. We calibrated under-reporting using independent estimates of transmission intensity provided by 12 serological surveys performed in 11 countries. We calculated local numbers of YF infections, deaths and disability-adjusted life years (DALYs) lost based on estimated transmission intensity while accounting for time-varying vaccination coverage. We estimated vaccine demand and impact of future preventive mass vaccination campaigns (PMVCs) according to various vaccination scenarios.

Vaccination activities conducted in Africa between 2005 and 2017 were estimated to prevent from 3.3 (95% CI 1.2-7.7) to 6.1 (95% CI 2.4-13.2) millions of deaths over the lifetime of vaccinees, representing extreme scenarios of all transmission due to the sylvatic or the urban cycle, respectively. By prioritizing provinces based on the risk of urban YF transmission, an average of 37.7 million annual doses for PMVCs over eight years would avert an estimated 9,900,000 (95% CI 7,000,000-13,400,000) infections and 480,000 (180,000-1,140,000) deaths over the lifetime of vaccinees, corresponding to 1.7 (0.7-4.1) deaths averted per 1,000 vaccine doses.

Limitations include substantial uncertainty in the estimates arising from the scarcity of reliable data from surveillance and serological surveys.

**Conclusions:** By estimating YF burden and vaccine impact over a range of spatial and temporal scales, while accounting for the specificity of urban transmission, our model can be used to inform the current EYE strategy.

---

Recent outbreaks in Angola, Nigeria and Brazil have shown that yellow fever (YF) remains a significant public health threat [1,2]. The epidemics of Zika and chikungunya in Latin America have also highlighted the risks of international spread of arboviruses. The spread of YF to Asia, where the virus has not yet been detected despite the presence of competent vectors, could have a major negative public health impact [3,4]. In response, in 2016, the World Health Organisation (WHO) adopted a strategy to Eliminate Yellow fever Epidemics (EYE) by 2026. This strategy aims to prevent sporadic cases sparking urban outbreaks, thus minimizing the risk of international spread [5]. The EYE strategy largely relies on, but is not limited to, scale-up of vaccination.

Vaccination activities considered in the EYE strategy consist of routine immunization of infants, preventive mass vaccination campaigns (PMVCs) that target all or most age groups, preventive catch-up campaigns targeting specific cohorts or unvaccinated sub-populations, and reactive campaigns in outbreak situations. Local assessment of YF transmission intensity is key for the prioritization of each of these vaccination activities; particularly since the supply of vaccine is limited as seen during the 2016 Angola outbreak [6,7]. Vaccine demand forecasts are critical to shape vaccine production and ensure optimal vaccine allocation.

Previous mathematical models have assessed geographical heterogeneity in YF risk [8–10]. However, modelling is challenging because of the co-existence of different transmission cycles [11]. In the sylvatic cycle, tree-dwelling mosquitoes transmit the virus within the wildlife reservoir (non-human primates) and spill-over infection may occur for humans living or working in jungle habitats. In the urban cycle, the domestic mosquito *Aedes aegyptii* transmits the virus between humans. Population (‘herd’) immunity (whether via natural infection or vaccination) will be expected to modify transmission intensity within the urban cycle, but not in the sylvatic cycle where transmission intensity is driven by non-human primates and their interactions with human populations. Recently, two models quantifying the incidence of the disease in Africa or worldwide have been used to derive estimates for vaccination impact [9,10]. Both assumed a constant force of infection over time, thus disregarding possible herd effects that may arise in urban context due to changing population-level immunity.

Accelerated urbanization and increasing human population mobility might increase the contribution of the urban transmission cycle to the global YF burden. Urban cases have the potential to trigger explosive outbreaks that can place a substantial burden on health systems, as well as causing significant social and economic impacts. Both the large number of cases arising in urban outbreaks, and the higher connectivity of affected populations compared to the, typically, much more remote settings in which sylvatic transmission occurs, make international spread more likely. Thus, accounting for specific features of urban transmission will be useful to assess the risk of urban outbreaks and refine the estimates of vaccine impact.

In this paper, we have extended of a previously developed model [9] to account for specific features of inter-human transmission. We then compare this model to the previous version (which focussed on sylvatic transmission) and update previous estimates of YF burden and vaccine impact, accounting for indirect effects. Lastly, based on local estimates of the potential for inter-human urban transmission, we propose different scenarios of PMVCs that could be considered for the EYE strategy, evaluating vaccine demand and impact in terms of infections and mortality averted.

## Material and methods

The Yellow Fever burden model [9] was developed to estimate YF disease burden and vaccine impact across 34 African countries at high or moderate risk for YF [5]. The original model was developed assuming that the locally-fitted force of infection (the annual probability of infection for a susceptible individual), *λ* was constant in time. Here, we present an alternative version of the model parametrised by the locally-fitted basic reproduction number *R_0_*, which allows the resulting force of infection to vary dynamically as population-level immunity changes over time.

### Model overview

A complete description of the model is available in S1 Appendix. Briefly, both model variants include a generalised linear model (GLM) fitted to the presence or absence of any reported YF event between 1984 and 2013 at the first sub-national administrative level (hereafter called province), using various environmental and demographic covariates. We defined a reported YF event as either outbreak reports published by WHO or laboratory-confirmed cases reported in a YF surveillance database managed by WHO-AFRO, to which 21 countries contribute.

The GLM provides estimates of the probability of any YF report across the endemic zone over the 30-year period considered. In a second step, we describe this probability of a report as dependent on the number of infections and on the unknown rate of under-reporting, which was calibrated to independent estimates of transmission intensity, obtained by fitting 12 serological surveys performed in 11 African countries.

The two model versions differ in the way they fit serological data. In the static version of the model (thereafter termed “FOI model”,) a constant, age-independent force of infection (FOI) *λ* is fitted to each serological survey. Alternatively, in the dynamical version of the model (thereafter termed “*R_0_* model”), a basic reproduction number *R_0_* is fitted, based on the classical SIR model framework under the assumption of endemic equilibrium transmission [12].

The GLM quantifies geographic variation of the relative risk of YF transmission across the continent while each serological survey yields an estimate of the absolute transmission intensity in its specific location. For each of the 31 survey locations we can therefore estimate the local level of under-ascertainment by tying GLM predictions to estimated values of *λ* or *R_0_* whilst accounting for time-dependent vaccination coverage [13]. By extrapolating this estimated level of under-ascertainment, we may infer the absolute transmission intensity across the continent from the GLM predictions.

The main difference in the dynamics of the FOI and *R*_0_ models lies in the way they respond to vaccination. In the FOI model, the susceptible population is reduced by vaccination but the per-capita risk of infection of remaining susceptible individuals remains unchanged, always resulting in a non-zero number of new infections with imperfect vaccination coverage. In contrast, the *R_0_* model responds non-linearly to vaccination coverage: if, for a specific year, vaccination activities happen to increase the immunized proportion of the population above the herd immunity threshold, *1 – 1/ R_0_* (also known as the Critical Vaccination Coverage, CVC), then no new infection will be expected in that year. In that case, the unvaccinated proportion of the population is indirectly but fully protected by herd immunity. Note, that when there is no history vaccination, both models agree.

### Model fitting and burden estimates

The model is fitted in a Bayesian framework using Markov chain Monte Carlo simulations. We assumed a prior distribution for vaccine efficacy centred at 97.5% (95% confidence intervals: 82.9% to 99.7%) [14], with no waning of immunity [15]. The models were fitted based on the best yearly estimates of vaccination coverage [13]. Posterior samples of parameters were used to compute medians and 95% credibility intervals (CI) of model parameters and burden estimates.

Both models predict spatiotemporally varying incidence of YF infections. We assumed that 12% (95% CI: 5% to 26%) of all infections develop severe disease and a case fatality ratio among severe cases of 47% (95% CI: 31% to 62%) [16] to translate infection incidence estimates into numbers of severe cases, deaths and disability-adjusted life years (DALYs) lost (S1 Appendix p12).

We further calculated vaccine impact of past vaccination activities by estimating the burden expected had these activities not taken place (S1 Appendix p12). We defined the lifetime impact of vaccination as the cumulative difference over the 2000-2100 time period in baseline burden estimates and those estimated for the counterfactual scenario of no vaccination. Such a time horizon ensures we capture vaccine impact over most of the lifetime of people vaccinated and those benefitting from the resulting herd immunity.

### Model validation

Additional data from three recent serological surveys conducted in the Democratic Republic of Congo, South Sudan and Chad in 2015 were withheld from the model fitting process and used for out-of-sample validation. Validation was conducted by comparing: i) transmission parameters estimated by fitting the age-seroprevalence profiles from those surveys to those predicted for the corresponding provinces in the YF burden model, ii) age-seroprevalence profiles directly observed in the surveys to those predicted by the YF burden model.

### Estimating vaccine demands and impact of large-scale vaccination campaigns

Lastly, we estimated the number of doses needed for, and the impact of, possible future PMVCs. As the EYE strategy aims to prevent outbreaks and international spread, we focused on the *R_0_* model which better captures urban transmission. We estimated the effective reproduction number *R_eff_* describing the transmission potential in a partially immune population across the endemic region as *R_eff_= R_0_(1-vc)*, where *vc* is the 2018 vaccine-acquired population-level immunity. Based on different *R_eff_* threshold values, four vaccination strategies were simulated in which provinces were eligible for PMVCs: i) *R_eff_*≥ 1.25, ii) *R_eff_*≥ 1.01, iii) *R_eff_*≥ 1.00, and iv) *R_eff_*≥0.85. For each strategy, the total target population was estimated. The corresponding number of vaccine doses was attributed to provinces based on a ranking of *R_eff_* values, so provinces with the highest *R_eff_* values were vaccinated first (2018) and those with the lowest *R_eff_* value were vaccinated last (2026). Vaccination campaigns were assumed to reach 90% of the population, regardless of age or previous vaccination status. For each strategy, the lifetime impact of PMVCs was estimated through comparison with a baseline scenario assuming no further reactive or preventive vaccination campaigns, but an annual 1%-increase in the coverage of routine immunization (capped at 90%) from their 2015 levels in countries where routine immunization already includes YF vaccine.

## Results

Based on the YF surveillance database and publicly-available reports, YF had been reported at least once over the 1984-2013 period in 160 of 479 provinces (Figure 1A). The GLM captured the presence/absence of YF well with an area under the ROC curve >0.9 for both model variants (Table 1 and S1 Figure).

**Figure 1:**
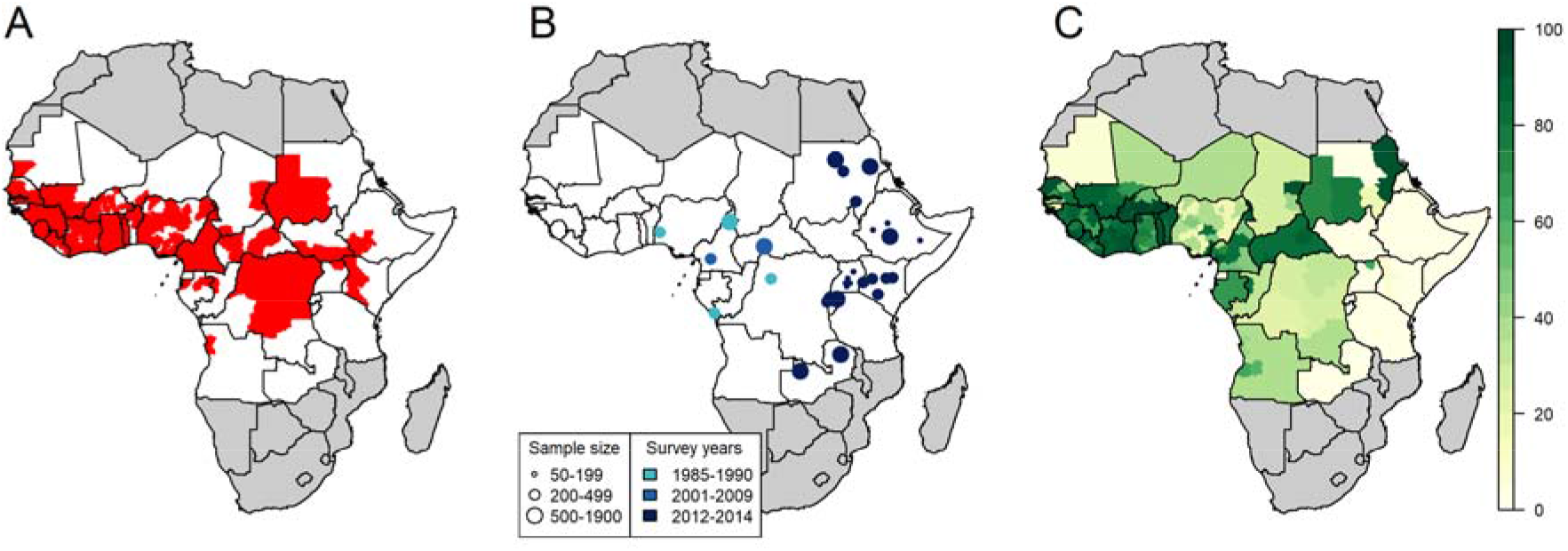
Input data of the model. A: presence (red) or absence (white) of any yellow fever report between 1984 and 2013. B: Location, sample size and study years 12 serological surveys covering 31 provinces. C: estimated population-level vaccination coverage for 2017.

**Table 1:** Parameter estimates and outcomes for both model variants.

|  | FOI model, median estimate (95%<br>Credibility Interval) | R<sub>0</sub> model, median estimate (95%<br>Credibility Interval) |
| --- | --- | --- |
| ***Model parameters*** |  |  |
| GLM Area under the Curve | 0.916 (0.909 - 0.921) | 0.916 (0.908 - 0.921) |
| Minimum per-infection<br>probability of detection | 3.6e-7 (2.1e-8 - 2.9e-6), Guinea-Bissau | 6.2e-7 (4.5e-8 - 3.6e-6), Guinea-Bissau |
| Maximum per-infection<br>probability of detection | 1.9e-5 (9.0e-6 - 3.8e-5), Central<br>African Republic | 3.0e-5 (1.8e-5 - 5.1e-5), Central<br>African Republic |
| Vaccine efficacy | 0.952 (0.749 - 0.993) | 0.942 (0.671 - 0.993) |
| ***Burden estimates*** |  |  |
| 1995 number of deaths | 110,000 (40,000 - 280,000) | 120,000 (50,000 - 320,000) |
| 2005 number of deaths | 130,000 (50,000 - 320,000) | 60,000 (20,000 - 210,000) |
| 2017 number of deaths | 110,000 (40,000 - 270,000) | 30,000 (4,000 - 120,000) |
| 2017 number of severe cases | 240,000 (90,000 - 620,000) | 70,000 (9,000 - 270,000) |
| 2017 total number of<br>infections | 2,190,000 (1,310,000 - 3,710,000) | 670,000 (100,000 - 1,790,000) |
| 2017 total number of DALYs<br>lost | 5,400,000 (1,900,000 - 13,600,000) | 1,700,000 (240,000 - 7,000,000) |
| ***Lifetime vaccine impact estimates\* of 2005-2017 PMVCs*** |  |  |
| Deaths prevented | 3,300,000 (1,200,000 - 7,700,000) | 6,100,000 (2,400,000 -<br>13,200,000) |
| DALY prevented | 145,800,000 (53,700,000 -<br>345,700,000) | 327,900,000 (133,000,000 -<br>699,300,000) |

DALYs: disability-adjusted life years; PMVCs: Preventive mass vaccination campaigns.
\*Lifetime vaccine impact is defined as the cumulative difference over the 2000-2100 time period in baseline burden estimates and those estimated had the 2005-2017 PMVCs not taken place. The 2000-2100 time horizon ensures to capture vaccine impact over most of the lifetime of people vaccinated and those benefitting from the resulting herd immunity.

We used the results of serological surveys conducted between 1985 and 2014 in 31 different locations (Figure 1B) to calibrate the models. Seroprevalence among unvaccinated participants ranged from 0.0% in northern Kenya in 2013 (95% CI: 0.0-0.9%) to 20.1% (15.0-26.5%) in South-West Nigeria in 1990. Population-level vaccination coverage estimates for 2017 ranged from 0.0% in various regions of Eastern Africa to 96.6% in several provinces of Burkina Faso (Figure 1C).

We estimated per-infection probabilities of detection by combining GLM predictions and results of serological surveys (Figure S2). These detection probabilities, which encompass all infections including those which are asymptomatic, were distributed across several order of magnitude around 10^−5^, but spatial heterogeneity was consistent between model variants (Table 1 and S2 Figure).

Both models consistently estimated the highest values of transmission intensity in terms of FOI and *R_0_* to be in West Africa and the lowest in Eastern Africa (Figure 2 and S3 Figure). Several provinces in Eastern and Central Africa had median *R_0_* estimates just above one – in reality these areas are likely not endemic for YF transmission, but the implicit assumption of endemic transmission means the *R_0_* model is unable to generate estimates of *R_0_<1*. Local CVC estimates were derived from *R_0_* values (S4 Figure). For instance, in Côte d’Ivoire, CVC estimates ranged between 49% and 98% across provinces. Both model versions estimated vaccine efficacy at 0.95, with lower bound of the 95% credibility interval surrounding 0.70 (Table 1).

**Figure 2:**
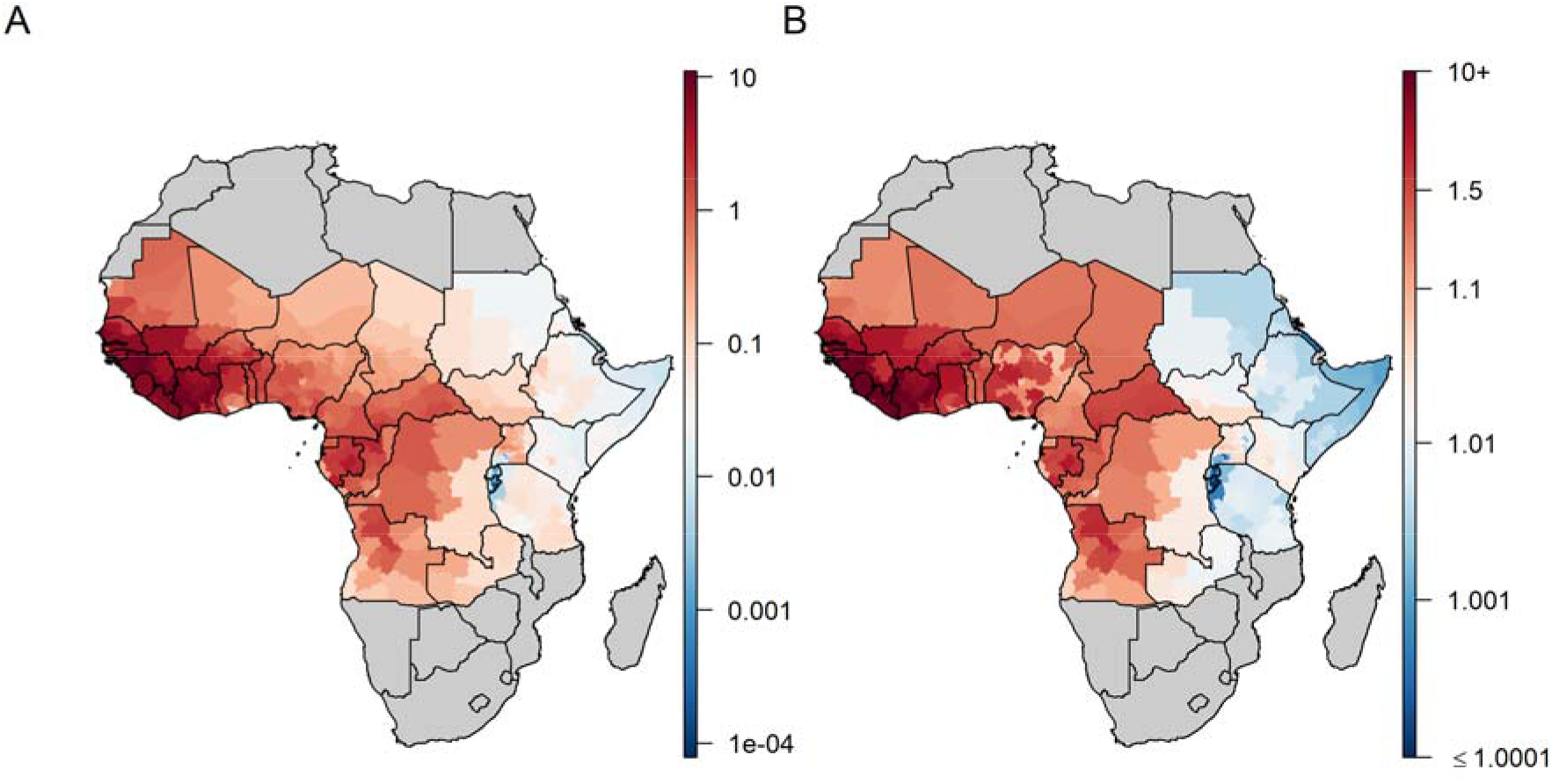
Estimates of transmission intensity across both model variants. A: median estimates of the force of infection (FOI), in %; B: median estimates of R_0_.

Results of the out-of-sample validation are presented in Figure 3 and S2 Appendix. Although uncertainty in the predictions were substantial, both model variants successfully reproduced heterogeneity in transmission intensity across provinces, particularly in South Sudan.

**Figure 3:**
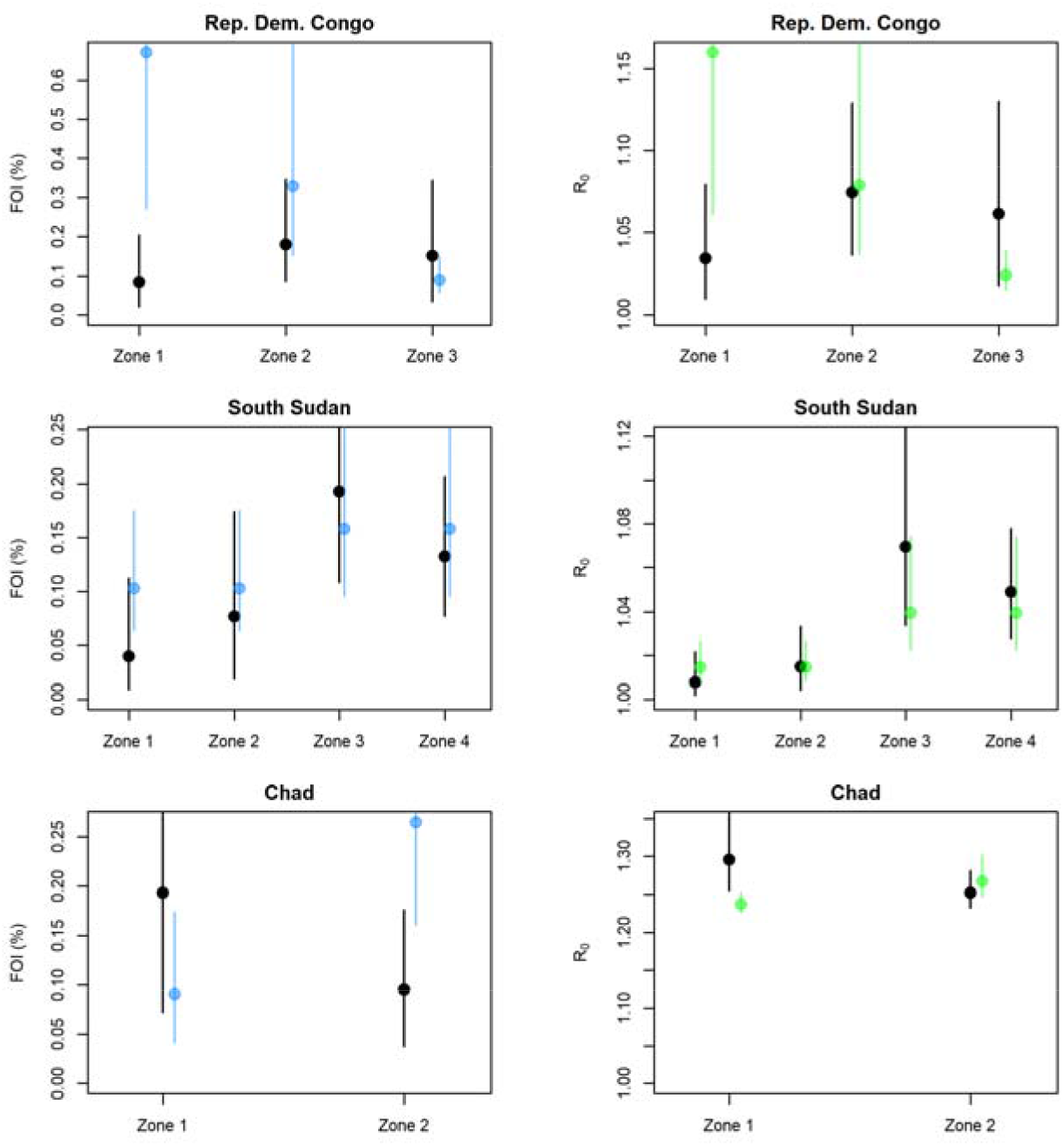
Comparison of directly estimated versus model-predicted transmission parameters for three external serological studies. Black: direct estimate; blue: prediction of the FOI model; green: prediction of the R_0_ model. Lines show 95% credibility intervals.

Estimates of the YF-specific burden are presented in Table 1. We estimate 107,000 deaths (37,000–272,000) occurred in 2017 according to the FOI model and 30,000 (4,000–123,000) according to the *R_0_* model. As expected from its non-linear response to vaccination, the *R_0_* model was more sensitive to variations in vaccination coverage and predicted a higher burden than the FOI model before the implementation of mass vaccination campaigns, but a lower burden thereafter (S5 Figure). Of note, with no or low-level of vaccination coverage as in Eastern Africa, both model versions exhibited similar temporal trends (S6 Figure). Province-based burden estimates are presented in Figure 4. While the assumption of endemic equilibrium forced the FOI model to predict a non-zero force of infection, large parts of the endemic zone exhibited very low incidence rates (notably in Rwanda and Burundi). In contrast, the *R_0_* model predicted a zero incidence in all provinces where the vaccination coverage exceeded the local CVC. Large regions of Eastern Africa, that did not benefit from any vaccination activities, still exhibited small incidence rates.

**Figure 4:**
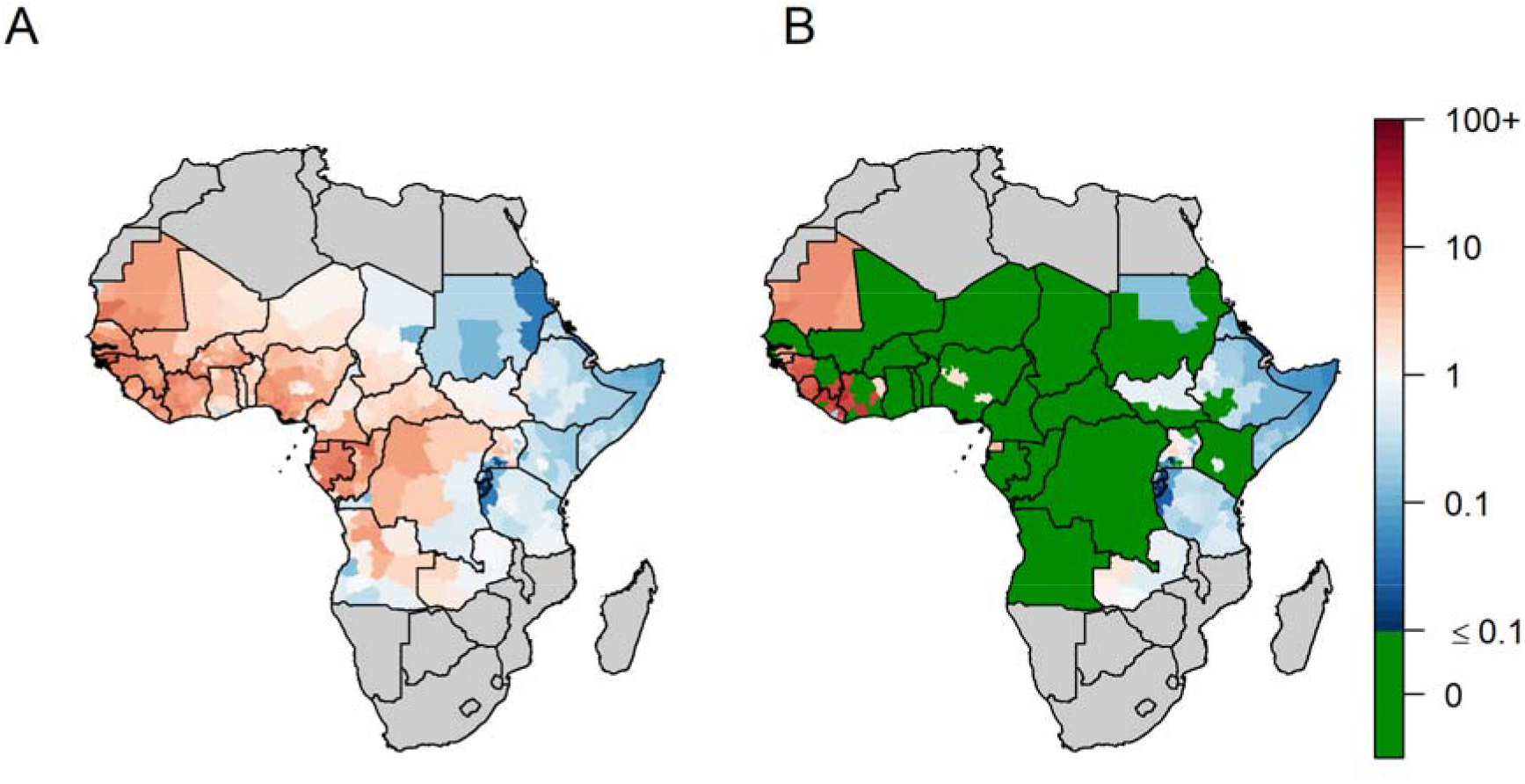
Estimated 2017 incidence of severe Yellow Fever infection per 100,000 persons across 34 African countries. A: FOI model, B: R_0_ model. For the R_0_ model, provinces with no incidence are those for which the estimated vaccination coverage in 2017 was larger than the Critical Vaccination Coverage implied by R_0_.

According to the FOI model, PMVCs conducted between 2006 and 2017 under the Yellow Fever Initiative [17] were estimated to have already prevented 700,000 (200,000-1,600,000) deaths as of 2017 and will prevent a further 2,600,000 (1,000,000-6,100,000) deaths over the lifetime of the vaccinees. According to the *R_0_* model, 2,600,000 (1,000,000-6,000,000) deaths have been prevented as of 2017 and 3,400,000 (1,300,000-7,400,000) future deaths will be prevented by those campaigns.

Figure 5 displays estimates of *R_eff_* in 2018 together with the categorisation of provinces for several PMVCs strategies. Due to low >1 *R_0_* values (by assumption) and non-existent vaccination coverage, large regions of Eastern Africa had estimates of 1.00<*R_eff_*<1.01. We estimated that an average annual number of 37.7 million doses would be sufficient to vaccinate all provinces with *R_eff_*≥1.01 over the 2018-2026 period. Such a strategy would prevent 9,900,000 (7,000,000-13,400,000) infections and 480,000 (180,000-1,140,000) deaths over the lifetime of the vaccinees, corresponding to 1.7 (0.7-4.1) deaths averted per 1,000 vaccine doses (Table 2). Broader strategies would result in larger number of deaths prevented at the cost of lower impact per doses.

**Figure 5:**
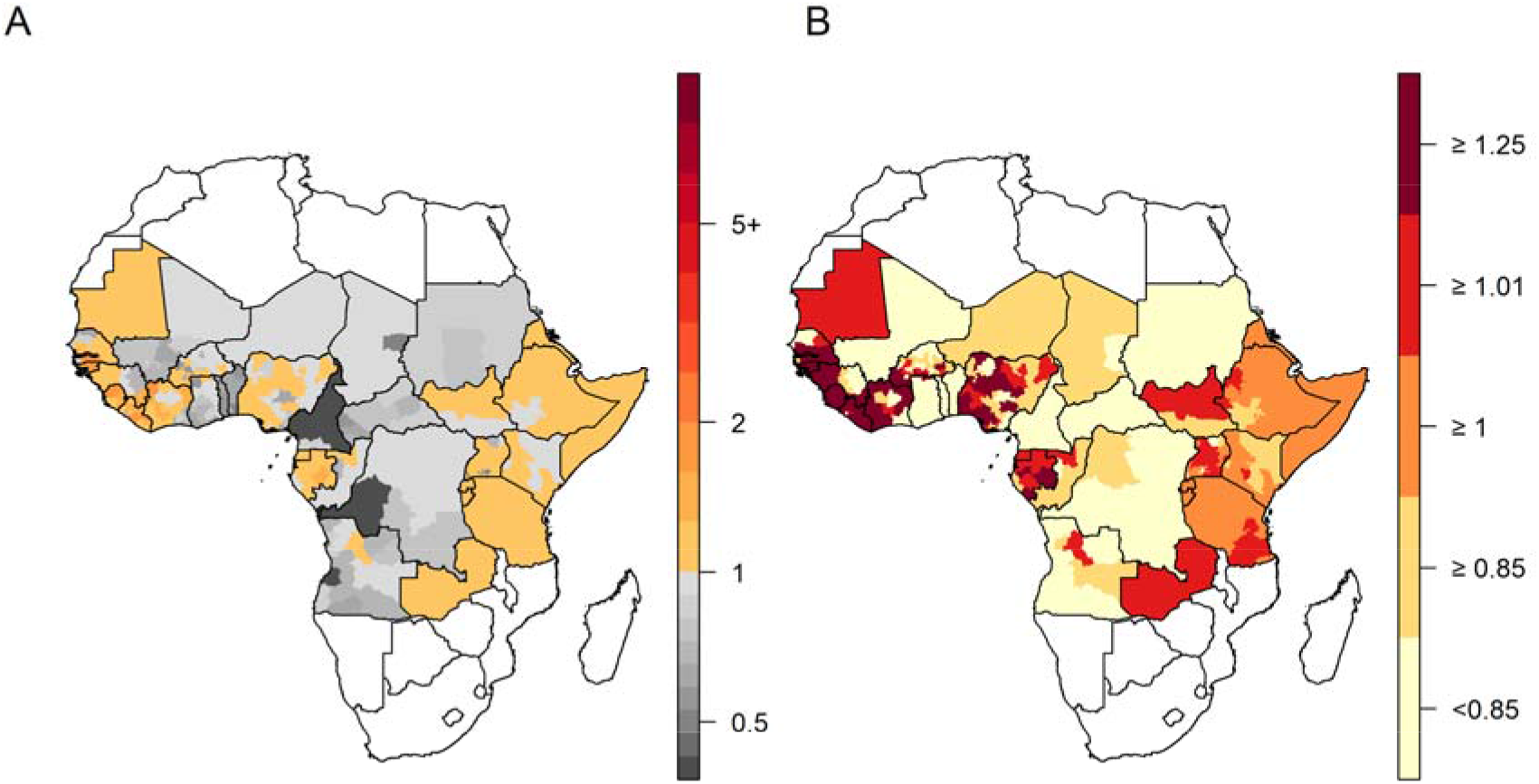
Estimates of the effective reproductive number (R_eff_) and categorization for different preventive mass vaccination (PMVCs) strategies. A: estimates of the effective reproductive number (R_eff_) based on 2018 vaccination coverage estimates; B: categorization for vaccination strategies. Each vaccination strategy includes all provinces above the given threshold (ie the strategy corresponding in vaccinating provinces with R_eff_≥ 1.01 consists in vaccinating provinces coloured in bright and dark red). Light yellow provinces (R_eff_<0.85) are not considered for PMVCs under any strategy.

**Table 2:** Impact estimates of different preventive mass vaccination (PMVCs) strategies. Infections and deaths prevented are calculated as the difference in the cumulative burden over the 2018-2100 time period between a scenario corresponding to the considered strategy and a baseline scenario assuming no future PMVCs. Infections include asymptomatic, mild and severe infections.

| EYE strategy | Total target population (millions) | Average number of doses per year | Infections prevented | Deaths prevented | Deaths prevented per 1,000 vaccine doses | % of deaths prevented |
| --- | --- | --- | --- | --- | --- | --- |
| $R_{eff} \geq 1.25$ | 143.9 | 18.0 | 1,378,000 (545,000-3,693,000) | 66,000 (16,000-235,000) | 0.5 (0.1-1.6) | 3% (1-7%) |
| $R_{eff} \geq 1.01$ | 277.9 | 37.7 | 9,888,000 (6,957,000-13,383,000) | 481,000 (182,000-1,144,000) | 1.7 (0.7-4.1) | 19% (8-46%) |
| $R_{eff} \geq 1.00$ | 505.3 | 63.2 | 14,244,000 (10,317,000-19,746,000) | 693,000 (270,000-1,607,000) | 1.4 (0.5-3.2) | 28% (11-73%) |
| $R_{eff} \geq 0.85$ | 644.4 | 83.1 | 15,141,000 (10,927,000-20,981,000) | 736,000 (288,000-1,716,000) | 1.1 (0.4-2.6) | 29% (12-77%) |

## Discussion

By combining a regression model fitted to the reported local presence of YF with serological data collected in 11 countries, this modelling study presents updated estimates of YF burden across 34 African countries in terms of number of infections, severe cases, deaths, and DALYs. We also estimated that PMVCs conducted since 2006 will prevent a total of 3 to 6 million deaths over the lifetime of vaccinees, depending on whether we assume transmission is dominated by the sylvatic or the urban cycles, respectively. As both transmission cycles do contribute to the disease burden, the real impact of vaccination likely lies between these two extreme cases. Finally, we derived estimates of vaccine demand and impact of future PMVCs under a variety of targeted vaccination scenarios. We estimated that allocating a 37.7 million doses per year between 2018 and 2026 for PMVCs would prevent 9.89 (6.96-13.3) million YF infections and thus considerably reduce the risk of international spread.

The model we developed combines a risk-map derived from occurrence data with transmission intensity estimates in order to infer burden. Such an approach also provides valuable results for other diseases, such as dengue [18]. We estimated approximately 200,000 annual YF severe cases occurred before 2005 (S5 Figure), that is before the implementation of PMVCs in West Africa. These results are consistent with previous estimates suggested over the recent decades [19,20], while providing better insight about spatial and temporal heterogeneity in the burden of the disease, especially for the post-PMVCs period. Uncertainty in our results is substantial, covering more than one order of magnitude for some estimates. As our model integrates all existing data on YF that are currently available for Africa, such uncertainty reflects the current lack of accurate knowledge regarding YF incidence and epidemiology.

The two model variants we developed exhibited substantial differences in resulting estimates of YF burden and vaccine impact, especially for the recent period. By integrating herd effects as an important component of urban transmission, the impact of vaccination estimated in the *R_0_* model was roughly twice that estimated in the FOI model, which represents sylvatic transmission. Cases’ characteristics and environmental assessment suggested a sylvatic origin of the 2016 Uganda outbreak [21]. Similarly, the lack of a large number of urban cases indicates a primarily sylvatic source for the 2017-2018 Nigeria outbreak [2]. On the contrary, the 2016 YF outbreak in Angola provided evidence of a large contribution of the urban transmission cycle [22]. Better understanding the relative contribution of the different transmission routes may affect control strategies. While vaccination prevents both sylvatic and urban transmission, vector population control interventions appear relevant in urban settings but may not be feasible to protect populations exposed to sylvatic mosquitoes. A recent study tried to disentangle the contribution of each transmission cycle in Brazil [23]. However, this analysis relied on detailed spatiotemporal and genomic data not currently available for Africa. Assumptions on the global contribution of both transmission cycles have important effects on vaccine impact estimates, which in turn may impact prioritization across diseases by vaccine funders. Combining estimates of model variants that rely on different assumptions and structures in a way that can reliably inform decision making remains a challenge.

The models used to estimate the burden and subsequently the impact of vaccination are fitted to data of reported YF events over a 30-years period in order to smooth over the long and irregular cycles of epidemic activity sweeping across the continent, such as the re-emergence during 1985-95 [24]. These data show a pattern of near-ubiquitous presence in large swathes of West Africa, with more patchy presence in Central and East Africa, which in turn determine the gradient in transmission intensity seen in the model. Due to the very widespread YF presence in West Africa, the model lacks resolution for estimating the exact level of transmission intensity in this area. Under-reporting of cases and transmission intensity are estimated by fitting serological surveys data from Central and Eastern Africa. Unfortunately, high quality surveys from West Africa are lacking. Of note, our validation assessment revealed that the model produces reasonable predictions across settings exhibiting varying levels of seroprevalence (S2 Appendix). This may be seen as a case for the validity of our model across a range of transmission intensity levels.

While the FOI model variant assumes a constant local infection risk for those unvaccinated, the *R_0_* model variant relies on the assumption of endemic equilibrium, which allows long periods without transmission if population-level vaccination coverage exceeds the CVC and therefore suppresses transmission [25]. However, this assumption prevents the model from representing the growth of large pools of susceptible individuals in the population between epidemics. As such, neither model variant completely captures the epidemic character of the disease driven by rapid depletion of susceptible hosts during explosive outbreaks, slow host renewal in inter-epidemic periods and sporadic disease reintroductions. Instead, these models were designed to capture spatial heterogeneity in the burden and estimate long-term average vaccine impact.

Both model fitting and projections are based on vaccine-coverage estimates derived from a demographic model combining different sources of data regarding vaccination activities [13]. Concurrent estimates for vaccine coverage have been recently developed [26]. Although overall consistent with ours, these estimates appeared to disregard several recent large-scale campaigns in West and Central Africa [13]. Uncertainty in vaccination coverage is not directly accounted for in our model, but may be propagated in the efficacy parameter of the model. Indeed, the posterior distribution of vaccine efficacy is much wider than the range documented in a recent meta-analysis [14], especially for the *R_0_* variant which is expected to be more sensitive to vaccine coverage. Lastly, in the absence of more detailed data, vaccination activities were often assumed to be implemented homogeneously across space in particular for RI, glossing over existing heterogeneities in coverage. Recent efforts to document such heterogeneity, such as the YF EPI warning system developed by the WHO, are thus welcome [27].

Our model has several limitations but also some strengths compared with past work. We captured spatial heterogeneity in the disease using a statistical rather than a mechanistic approach. Our model does not explicitly represent the spatial distribution of vectors nor disease reservoirs unlike Shearer et al’s model [10]. However, the set of spatial predictors we used has been demonstrated to have good predictive power for both presence and seasonality of YF [28]. To our knowledge, our model is the only one calibrated against serological data that captures the whole spectrum of disease severity. Other YF burden estimates stem from models calibrated on reported cases [10,29], that may lead to underestimation due to under-reporting and misdiagnosis. For instance, the Global Burden of Disease estimates for 2016 around 5,000 YF-deaths in Africa, which is near to the lower bound of our *R_0_* model estimates. Lastly, the development of the *R_0_* variant of our model constitutes a novel approach to account for herd effects in the impact of YF vaccination. Our results highlight that heterogeneity in transmission intensity imply different vaccination threshold to be reached in order to prevent YF outbreaks. This can be used to locally adapt the empirical threshold of 80% vaccination coverage that is often considered sufficient for YF outbreak prevention [11].

Using local estimates of the potential for urban outbreaks for PMVCs prioritization, we explored several vaccination scenarios ranking from a parsimonious strategy focussed on high-risk areas to an ambitious large-scale approach targeting the vast majority of provinces within the at-risk region. Recent years have shown that efforts to control YF outbreaks were highly constrained by the stock of vaccine doses available for emergency situation [30]. Optimal vaccine allocation is also crucial in the context of preventive vaccination activities. By contrasting the impact of several scenarios with the corresponding vaccine demand, and also by suggesting a sequence of implementation within each scenario, this study helps in the search for optimal vaccine allocation. We do acknowledge however that a realistic strategy will need to take into account further factors such as operational or political concerns.

Patterns of YF transmission seem to have changed in the recent years toward shorter transition from the sylvatic cycle to urban inter-human transmission [5]. Renewed disease control strategies thus need to account for this change in implementation of control measures. This study addressed this challenge by developing a vaccine impact modelling framework accounting for urban YF transmission. This work has contributed to a risk assessment conducted as part of the EYE strategy in order to prioritize countries for PMVCs [31].

## Supporting information

**S1 Appendix:** Model description.

**S2 Appendix:** Model validation.

**S1 Figure:** Observed versus predicted presence/absence of any yellow fever reported event between 1984 and 2013.

**S2 Figure:** Country-specific per-infection probability of detection across both model variants.

**S3 Figure:** Variability in the estimates of transmission intensity across model variants.

**S4 Figure:** Estimated median critical vaccination coverage (A, %) and interquartile range in the critical vaccination coverage (B, %).

**S5 Figure:** Comparison of yellow fever burden estimates over time between both model versions across the whole endemic region (34 countries).

**S6 Figure:** Comparison of yellow fever burden estimates over time between both model versions across sub-regions.

## Acknowledgement

We thank Olivier Ronveau and William Perea for useful comments on this work. We are also thankful to Sergio Yactayo and Messeret Shibeshi for granting access to epidemiological data. This work was carried out as part of the Vaccine Impact Modelling Consortium (www.vaccineimpact.org), but the views expressed are those of the authors and not necessarily those of the Consortium or its funders. The funders were given the opportunity to review this paper prior to publication, but the final decision on the content of the publication was taken by the authors.

## Abbreviations

CI: credibility interval
CVC: critical vaccination coverage
DALYs: disability-adjusted life years
EYE: Eliminate Yellow fever Epidemics
FOI: force of infection
GLM: generalized linear model
PMVCs: preventive mass vaccination campaigns
WHO: World Health Organization
YF: yellow fever

